# Formate reduces ischemic injury in the male heart by increasing protein *S*-nitrosation

**DOI:** 10.64898/2026.02.24.707768

**Authors:** Haley Garbus-Grant, Raihan Kabir, Obialunanma V. Ebenebe, Priyanka Patel, Deepthi Ashok, Diego Quiroga, Samarjit Das, Brian O’Rourke, Mark Crabtree, Mark J. Kohr

**Author notes:** **CORRESPONDING AUTHOR:** Mark J. Kohr, PhD, FAHA, FCVS, Department of Environmental Health and Engineering, Johns Hopkins Bloomberg School of Public Health, 615 N. Wolfe Street, Room E7616, Baltimore, MD 21205.

## Abstract

Ischemic heart disease is a leading cause of death for both men and women in the United States. We and others have demonstrated that nitric oxide (NO) signaling and associated protein *S*-nitrosation (SNO) play a key role in reducing ischemic injury in the heart. We also find that while females typically exhibit endogenous protection from ischemic injury, this protection is abrogated with the loss of the formate-generating enzyme alcohol dehydrogenase 5 (ADH5), but formate supplementation provided a rescue. Here, we investigate the cardioprotective efficacy of formate in male hearts. Hearts were Langendorff-perfused and subjected to ischemia/reperfusion (I/R) injury with and without formate. Formate-mediated protection was also examined using an *in vitro* model of coverslip-induced ischemic injury to identify molecular underpinnings. We found that formate yields protection from I/R injury in *ex vivo a*nd *in vitro* models by increasing post-ischemic protein SNO levels, while NO synthase inhibition blocked this formate-mediated increase in protein SNO *in vitro*, and attenuated protection from I/R injury *ex vivo.* Moreover, post-ischemic levels of tetrahydrobiopterin (BH_4_), a cofactor necessary for NOS function, were preserved in formate-treated hearts. Furthermore, inhibition of dihydrofolate reductase (DHFR), a one-carbon enzyme critical for BH_4_ recycling, blunted formate-mediated protection *ex vivo*. Collectively, our findings suggest that formate is a potent cardioprotective agent that confers protection by preserving post-ischemic BH_4_ levels, and enhancing protein SNO levels through a NOS-dependent mechanism. These findings have significant implications for the clinical prevention and treatment of ischemic heart disease in males.

## INTRODUCTION

Ischemic heart disease is a leading cause of morbidity and mortality among men and women in the US, affecting ∼5% of the US population.^1, 2^ Timely reperfusion represents the only effective clinical strategy for salvaging ischemic myocardium in both men and women, but significant injury results from reperfusion, due in part to the formation of reactive oxygen species (ROS) and the disruption of cellular redox balance.^3^ Cardioprotective strategies hold promise for combating ischemia-reperfusion (IR) injury in preclinical models by reducing infarct size and maintaining redox homeostasis.^4, 5^ Redox balance is maintained, in part via antioxidants, but nitric oxide (NO) also plays a role. In particular, NO has emerged as an important signaling molecule in the heart, yielding protective benefits through multiple pathways.^5^ NO is enzymatically produced via NO synthase (NOS) with tetrahydrobiopterin (BH_4_) serving as an essential co-factor. NO commonly signals via protein *S*-nitrosation (SNO), a posttranslational modification that we and others have shown to act as an important component of cardioprotection in preclinical models,^6–14^ but clinical translation remains elusive.^15–19^

Our recent findings reveal that NO-derived protein SNO represents a critical stress response to ischemia that is essential for maintaining redox homeostasis, as we have demonstrated that cardioprotective interventions that increase protein SNO, strongly correlate with reduced oxidative stress and ischemic injury, particularly in males.^6–14^ Moreover, in studies of both male and female hearts, we found that compared to males, female hearts exhibited higher baseline protein SNO levels and, associated with this, better maintenance of redox homeostasis and reduced susceptibility to myocardial I/R injury.^12, 20–22^ We also found that, compared to males, female hearts exhibited greater activity of alcohol dehydrogenase 5 (ADH5), a highly conserved dual function enzyme that acts as a reductase, regulating protein SNO levels, and a dehydrogenase, mediating the metabolism of formaldehyde to formate.^12^ Importantly, the loss of ADH5 blocked female-specific cardioprotection by worsening ischemic injury in female hearts.^21^ However, activation of mitochondrial aldehyde dehydrogenase 2 (ALDH2), which also metabolizes formaldehyde to formate, rescued female ADH5^-/-^ hearts.^21^ Further supporting a protective role for formate in the female heart, we recently found that exogenous formate restores protection against I/R injury in female hearts lacking ADH5.^23^ However, the mechanism underlying formate-mediated cardioprotection is not known, and whether it provides I/R injury protection in male hearts has not been explored.

In the current study, we uncover a protective role for formate in male hearts, and explore the underlying mechanism(s) of protection. We found that formate substantially reduced infarct size in Langendorff-perfused male hearts and reduced injury in an *in vitro* model using neonatal mouse cardiomyocytes (NVMMs) subjected to coverslip-induced ischemia. Formate-mediated protection was associated with an increase in protein SNO and preservation of BH_4_ levels after ischemia. Moreover, protection was alleviated with either non-specific NOS or dihydrofolate reductase (DHFR) inhibition, suggesting that formate may confer protection in male hearts through NOS- and DHFR-dependent signaling mechanisms.

## MATERIALS AND METHODS

### Animals

This investigation conforms to the Guide for the Care and Use of Laboratory Animals published by the National Institutes of Health (NIH; Publication No. 85-23, Revised 2011), which was approved by the Institutional Animal Care and Use Committee of The Johns Hopkins University. Male and female wild-type (WT, C57BL/6J) mice were purchased from the Jackson Laboratory (Bar Harbor, ME) and used between 15 and 32 weeks of age. All mice were housed in a pathogen-free environment and maintained on standard lab chow. Experiments were conducted at least one week after arriving on-site at our animal facility.

### Langendorff Heart Perfusion

Mice were anesthetized with a mixture of ketamine (90 mg/kg; Hofspira, Hempstead, NY) and xylazine (10 mg/kg; Sigma-Aldrich, St. Louis, MO) via intraperitoneal injection and were anticoagulated with heparin (Fresenius Kabi, Lake Zurich, IL). Hearts were excised and cannulated on a Langendorff apparatus, and perfused retrogradely with Krebs-Henseleit buffer (KHB) (95% O_2_, 5% CO_2_, pH 7.4) under constant pressure (100 cmH_2_O) and temperature (37°C) as previously described.^12, 20, 21, 23–26^ KHB consisted of (in mmol/L): NaCl (120), KCl (4.7), KH_2_PO_4_ (1.2), NaHCO_3_ (25), MgSO_4_ (1.2), d-glucose (11), and CaCl_2_ (1.75). For the ALDH2 activity assay and western blots, hearts were perfused for 5 mins and then snap-frozen in liquid nitrogen and stored at −80°C until use. Otherwise, hearts were subjected to the I/R injury protocol described below.

### Ex Vivo Ischemia/Reperfusion Injury Protocol

Hearts were subjected to I/R injury as previously described;^12, 20, 21, 23–26^ all procedures were performed in the dark to protect against light-induced cleavage of protein SNO. Briefly, hearts were cannulated on the Langendorff apparatus as described above. For functional assessments, a polyethylene balloon was inserted into the left ventricle to measure left ventricular developed pressure (LVDP) and heart rate using a PowerLab system (AD Instruments, Dunedin, New Zealand). All hearts undergoing I/R injury were initially perfused with control KHB for a 10-minute equilibration period. After equilibration, hearts were perfused under one of the following conditions: 1) control KHB, 2) KHB with 30 μmol/L sodium formate (Sigma-Aldrich), 3) KHB with 10 μmol/L Nω-Nitro-L-arginine methyl ester hydrochloride (L-NAME, Sigma-Aldrich), 4) KHB with 30 μmol/L sodium formate and 10 μmol/L L-NAME, or 5) KHB with 30 μmol/L sodium formate and 10 μmol/L methotrexate (Sigma-Aldrich). Hearts were perfused with respective treatments for 20 mins prior to ischemia, at which point the flow of buffer was halted for 25 mins to induce global, normothermic ischemia. After ischemia, the flow of buffer was restored, and hearts were reperfused with each respective treatment. Hearts undergoing molecular analysis were sectioned into halves or quarters and snap frozen in liquid nitrogen after 2 or 15 mins of reperfusion. Hearts evaluated for infarct size were reperfused for 90 mins. For infarct size analysis, hearts were perfused and incubated with tetrazolium trichloride (TTC, Sigma-Aldrich) dissolved in KHB buffer at 37°C for 20 mins followed by fixation in formalin. Non-ischemic hearts used for molecular analysis were perfused with respective treatments on the Langendorff apparatus for the same amount of time as hearts subjected to I/R injury.

### Infarct Analysis

After formalin fixation for a minimum of 24 hrs, hearts were cut into 1-2 mm cross-sections and imaged using a dissecting scope (Leica, Teaneck, NJ). Infarct size was then assessed using ImageJ software (NIH, Bethesda, MD) as previously described.^12, 20, 21, 23–26^ Infarct size is expressed as a percentage of the total area of the ventricle. All samples were blinded prior to sectioning and infarct analysis.

### Neonatal Mouse Ventricular Cardiomyocyte Isolation

Neonatal mouse ventricular cardiomyocytes (NMVMs) were isolated from postnatal day zero to three pups using the MACS cell separation kit (130-100-825, 130-098-373; Miltenyi Biotec, Bergisch Gladbach, Germany), as previously described.^27, 28^ Pups were decapitated, and hearts were excised and placed in sterile Dulbecco’s phosphate-buffered saline on ice (Gibco). Hearts were then minced into small pieces and digested at 37°C for 15 mins using the enzymes provided with the kit. Neonatal hearts were then mechanically digested using the m_neoheart_01 program on the gentleMACS dissociator (Miltenyi Biotec) for a total of 3 incubations and 3 runs. Enzymatic digestion was halted via addition of cold isolation media (Medium-199 supplemented with 25 mM HEPES, 2 μg/mL vitamin B12, 50 U/mL P/S, 1× nonessential 286 amino acids, and 10% FBS). Red blood cells were lysed with ACK lysis buffer (Gibco). Noncardiomyocytes were labeled with magnetic beads (Miltenyi Biotec). The total cell suspension was passed through a magnetic field to obtain a cardiomyocyte-rich cell suspension. Cells were seeded at a density of 1×10^6^ on fibronectin-coated glass-bottom dishes 35-mm (D = 20 mm) (NEST, Jiangsu, China) in isolation media. The next day, the media was changed to maintenance media (Medium-199 supplemented with 25 mM HEPES, 2 μg/mL vitamin B12, 50 U/mL P/S, 1× nonessential 286 amino acids, and 2% FBS).

### Coverslip-induced Ischemia/Reperfusion Injury Protocol

NMVMs were seeded on fibronectin-coated dishes and allowed to attach for 24 hrs and then subjected to a model of coverslip-induced I/R injury as described previously at 37°C.^27, 29, 30^ Cells were pretreated for one hour with fresh maintenance media (2% FBS, as described above) containing their respective treatment, 30 μmol/L sodium formate (Sigma-Aldrich), or 30 μmol/L sodium formate + 10 μmol/L L-NAME. After one hour of pretreatment, a glass coverslip was placed on the cells to induce ischemia for one hour,^30^ followed by the removal of the coverslip for a one-hour reperfusion period.^31^ At the completion of reperfusion, the media was collected, transferred to clean tubes, and snap-frozen in liquid nitrogen.

### Lactate Dehydrogenase Assay

Lactate dehydrogenase (LDH) activity was assessed using the CyQuant LDH Cytotoxicity Assay (ThermoFisher). Briefly, media from NMVMs subjected to coverslip induced I/R injury as described above was collected, and samples were assayed for LDH activity in triplicate using a 96-well plate. The reaction mixture was prepared per the manufacturer’s instruction, and the plate was incubated at 37°C for 30 mins and protected from light. The stop solution was added, and absorbance was read at 490 nm and 680 nm. Absorbance values at 680 nm (background) were subtracted from values at 490 nm. Each media type was also assayed in the absence of cells and subtracted from their respective experimental conditions to control for inherent LDH activity in serum. Absorbance values were normalized to non-treated I/R injury conditions in each isolation so that LDH release is expressed as a percentage of control.

### H9c2 Cell Culture

H9c2 cells were obtained from the American Type Culture Collection (ATCC, Manassas, VA) and grown in Dulbecco’s Modified Eagle Medium (DMEM) with 5% penicillin/streptomycin (P/S) and 10% fetal bovine serum (FBS). Cells were grown in 25 cm^2^ tissue culture flasks and treated with either control media, media with 10 μmol/L formate, or media with 10 μmol/L formate + 10 μmol/L L-NAME for 48 hrs once confluence was reached (∼2.8×10^6^ cells). Cells were then lysed in either sucrose buffer for the modified-biotin switch assay for protein SNO or in lysis buffer (Cell Signaling Technology) for western blot, as described above.

### Tissue Homogenization

For the ADH5 activity assay and western blot, hearts were homogenized in 1x lysis buffer (Cell Signaling Technology, Danvers, MA) with protease/phosphatase inhibitor cocktail and neocuproine. For the ALDH2 assay, hearts were homogenized in Dulbecco’s phosphate-buffered saline (Gibco, Carlsbad, CA) with protease/phosphatase inhibitor cocktail (Cell Signaling Technology) and neocuproine. For the biotin switch assay for protein SNO, heart quarters were homogenized in buffer containing (in mmol/L): sucrose (300), HEPES-NaOH pH 8.0 (250), EDTA (1), neocuproine (0.1), Triton-X 100 (0.5%), and an EDTA-free protease inhibitor tablet (Roche, Indianapolis, IN). Mechanical homogenization was performed using a Precellys Evolution 24 homogenizer (2*30s cycles, 0°C, 7200 RPM; Bertin Instruments, Montigy-le-Bretonneux, FR) with a hard tissue lysing kit. Homogenates were then assayed for protein concentration using a Bradford assay, and aliquots of tissue homogenates were stored at −80°C until use.

### ALDH2 Western Blot

Heart homogenate samples (30 μg) were separated (20 mins @ 75V; 70 mins @ 175 V) on a gradient Bis-Tris SDS-PAGE gel (4-12%, NuPAGE; Thermo Fisher). Gels were loaded with at least one molecular weight marker. Proteins were then transferred (90 mins, 220 mA, 30V) to a PVDF membrane (Thermo Fisher). Total protein was quantified using Ponceau S staining (Sigma-Aldrich). Membranes were then blocked in (5% wt/vol) powdered milk (BioRad, Hercules, CA). Membranes were then incubated with anti-ALDH2 (1:1000, Cell Signaling Technology, 18818S) overnight at 4°C in blocking solution. Membranes were then incubated for 1 hour at room temperature with the appropriate secondary anti-mouse antibody (1:5000; Cell Signaling Technology, 7076S). Chemiluminescence substrate (SuperSignal West Pico PLUS, Thermo Fisher) was added to the membrane and visualized using an iBright imager (Thermo Fisher). Protein expression was quantified using ImageJ software (National Institute of Health) and normalized to total protein.

### Modified Biotin Switch Assay for S-Nitrosation

This procedure was performed in the dark, as previously described.^8, 21, 32^ Briefly, heart homogenates were prepared in sucrose buffer as described above, and samples (100 μg protein) were diluted in HEN buffer: **H**EPES-NaOH pH 8.0 (250 mmol/L), **E**DTA (1 mmol/L), **N**eocuproine (0.1 mmol/L). Samples were then incubated with 2% SDS and 20 mM N-ethylmaleimide (NEM) at 50°C for 40 mins at 800 rpm. Following incubation, samples were acetone-precipitated and incubated at −20°C for 20 mins. Samples were then centrifuged for 10 mins at 10,000 x g at 4°C, and the supernatant was removed. The remaining protein pellet was then air dried and resuspended in HEN with 1% SDS using a motorized pestle. DyLight Maleimide (4 mmol/L, Pierce, Rockford, IL) was then added to each sample along with ascorbate for specificity (1 mol/L, 1:50 v/v), followed by incubation in the dark for 1 hour at room temperature. Following the incubation, samples were again acetone-precipitated and resuspended in lithium dodecyl sulfate (LDS) sample buffer (Thermo Fisher, Carlsbad, CA). Samples were then loaded onto a gradient Bis-Tris SDS-PAGE gel (4-12%, NuPAGE; Thermo Fisher) and run at 75 V for 20 mins, followed by an additional 100 mins at 150V. Gels were then visualized using an iBright imager (Thermo Fisher) in fluorescence mode.

### ALDH2 Activity Assay

Hearts were homogenized in Dulbecco’s phosphate-buffered saline (Gibco) with protease/phosphatase inhibitor cocktail (Cell Signaling Technology) and neocuproine as described above. After incubating for 20 mins on ice, samples were centrifuged at 16,000 g at 4°C for 20 mins. ALDH2 activity was then assessed using a commercial colorimetric kit (Abcam, Waltham, MA) per the manufacturer’s instruction. Activity was estimated by measuring the conversion of NAD^+^ to NADH at an absorbance of 450 nm every minute for 2 hrs.

### ADH5 Activity Assay

ADH5 activity was assessed in whole heart homogenates as previously described.^12, 20, 21, 23, 33^ Heart homogenates (100 μg) were diluted in assay buffer containing (in mmol/L): Tris-HCl pH 8.0 (20), EDTA (0.5), neocuproine (0.5) with 0.1% NP-40 and protease/phosphatase inhibitor cocktail (Cell Signaling Technology). NADH (200 μmol/L, Sigma-Aldrich) and GSNO (400 μmol/L, Sigma-Aldrich) were then added to initiate the reaction. NADH consumption was monitored via absorbance at 340 nm over 30 mins at 25°C. Activity was then determined based on the rate of NADH consumption.

### BH_4_ and BH_2_ Assessments

BH_4_, BH_2,_ and total biopterin (B) in heart homogenates were determined by high performance liquid chromatography (HPLC) followed by electrochemical and fluorescence detection, as previously described.^34^ Briefly, hearts were resuspended in 300 μL of phosphate buffered saline (50 mmol/L; pH 7.4), containing dithioerythritol (DTE; 1 mmol/L) and EDTA (100 μmol/L), and homogenized. After centrifugation (15 mins at 13,000 rpm and 4 °C), samples were transferred to new, cooled microtubes and precipitated with cold phosphoric acid (1 mol/L), trichloroacetic acid (2 mol/L), and DTE (1 mmol/L). Samples were vigorously mixed and then centrifuged for 15 mins at 13,000 rpm and 4°C, as described previously.^34^ Heart samples were then injected onto an isocratic HPLC system and quantified using sequential electrochemical (Coulochem III; ESA Inc., MA) and fluorescence (Jasco, UK) detection. HPLC separation was performed using a 250-mm, ACE C-18 column (Hichrom, UK) and mobile phase comprising sodium acetate (50 mmol/L), citric acid (5 mmol/L), EDTA (48 μmol/L), and DTE (160 μmol/L) (pH 5.2) at a flow rate of 1.3 ml/min. All reagents were ultrapure electrochemical HLPC grade. Background currents of + 500 and − 50 μA were used for the detection of BH_4_ on electrochemical cells E1 and E2, respectively. BH_2_ and biopterin were measured using a Jasco FP2020 fluorescence detector. Quantification of BH_4_, BH_2_, and biopterin was done by comparison with authentic external standards and normalized to sample protein content. B levels are expressed as the sum of detectable BH_4_, BH_2_, and biopterin.

### Statistics

Data were analyzed using GraphPad Prism (La Jolla, CA). Results are expressed as the mean +/- standard error of the mean. Outliers were identified using the ROUT method (Q = 1%), and normality of distributions was assessed using a Shapiro-Wilk test, as described previously.^23^ Significance was determined using a Students t-test or a one-way ANOVA with Tukey’s multiple comparisons for normally-distributed data. For non-normally distributed data, significance was determined using the Mann-Whitney rank sums test for two groups or a Kruskal-Wallis test with Dunn’s multiple comparisons correction. Significance was set at p<0.05. For enzyme assays, the linearity of the data was determined by a best-fit line. Linear regression analysis was used for statistical comparison of linear data, (i.e. enzyme activity assays). Curve differences were evaluated with 95% confidence intervals.

## RESULTS

### ALDH2 activity is decreased in male hearts

A prior study showed that female rat hearts have higher ALDH2 activity compared to males,^35^ so we initially sought to determine whether this difference in ALDH2 activity was also present in male and female mouse hearts. We quantified ALDH2 expression and found that while protein expression levels showed a trend towards significantly lower ALDH2 levels in females hearts vs. males (Fig. 1A), ALDH2 activity was significantly less in male hearts compared to females (Fig. 1B). Taken together with our previous findings on ADH5 activity,^12, 20, 21^ male mouse hearts show reduced ADH5 and ALDH2 activity compared to females.

**Figure 1.**
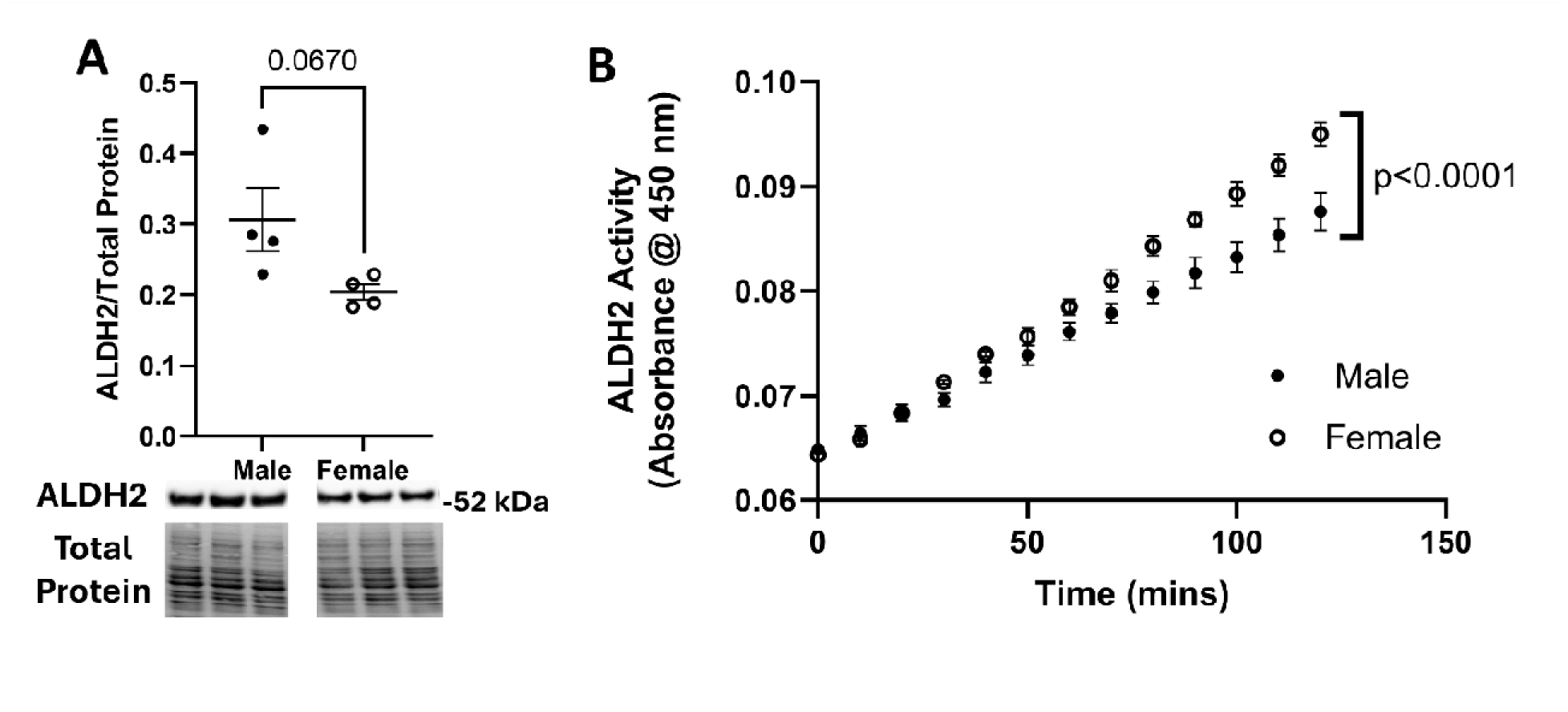
ALDH2 activity is higher in female hearts. Male and female hearts were Langendorff-perfused under control conditions. **(A)** ALDH2 protein expression via SDS-PAGE and western blot normalized to total protein measured using Ponceau S stain (p = 0.0670; n = 4 hearts/group) and **(B)** activity via NADH-linked assay in male and female hearts (p<0.0001; n = 4 hearts/group).

### Formate protects male hearts from ex vivo I/R injury

Exogenous formate administration reduced ischemic injury in female hearts lacking ADH5 in our previous study.^23^ Since formate is produced via ALDH2- and ADH5-mediated formaldehyde metabolism, and male hearts show reduced ADH5^12, 20, 21^ and ALDH2 (Fig. 1) activity, we hypothesized that exogenous formate may yield protective benefits in male hearts subjected to I/R injury. Indeed, we found that the administration of 30 μmol/L formate prior to ischemia and at the onset of reperfusion, substantially reduced ischemic injury in male hearts, with male hearts exhibiting a significant decrease in infarct size (Fig. 2A-B) and a significant increase in functional recovery (Fig. 2C). Conversely, female hearts did not show a protective benefit with formate (Fig. 2A-C), which is consistent with our prior study.^23^ As such, our findings suggest that formate is highly effective at reducing ischemic injury in male hearts, but similar benefits were not observed in females.

**Figure 2.**
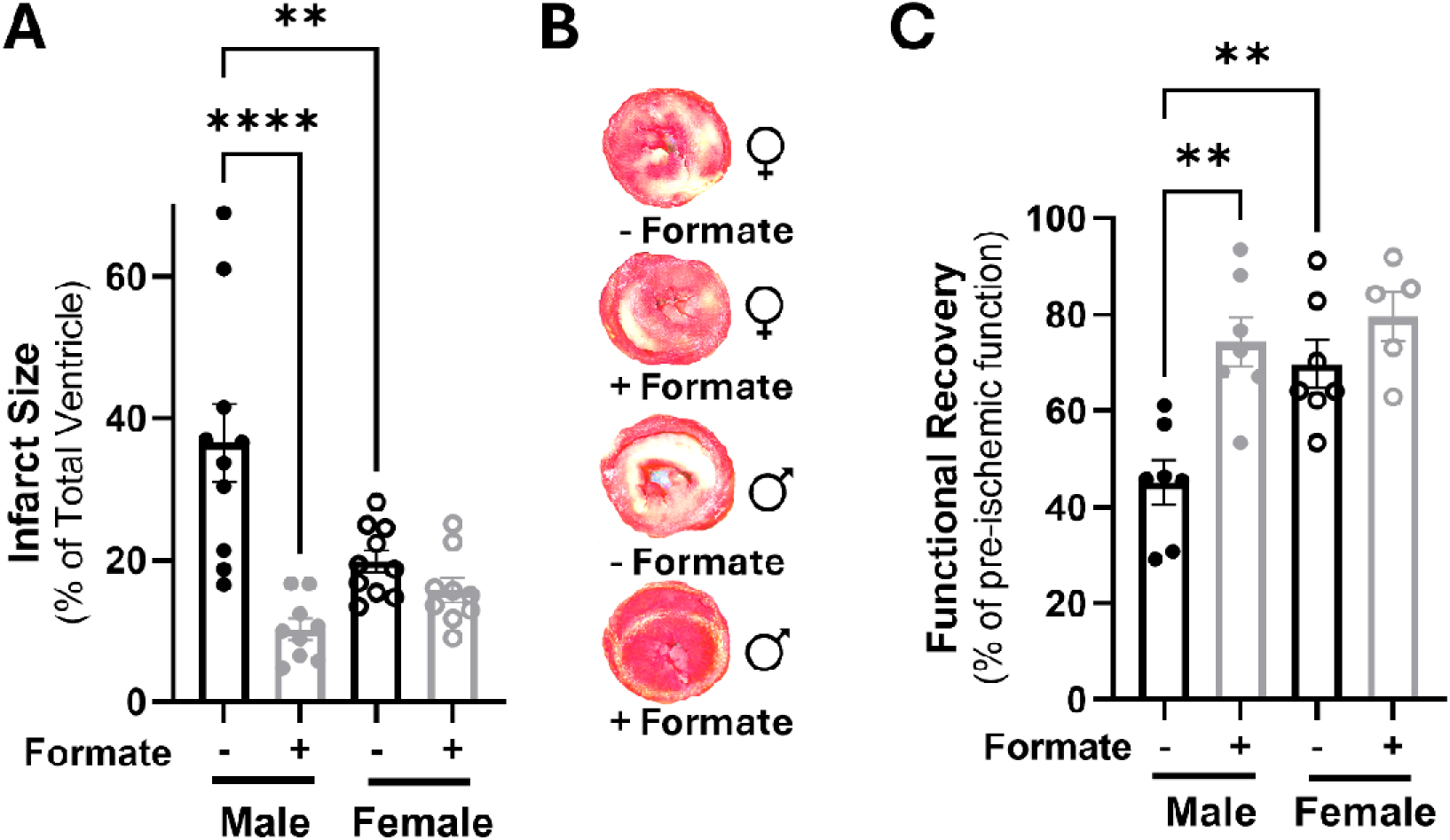
Exogenous formate reduces I/R injury in male hearts. Male and female hearts were Langendorff-perfused +/- 30 µmol/L formate and subjected to I/R injury. **(A)** Infarct size (****p<0.0001 vs. male + formate, **p<0.01 vs. female; n = 9-10 hearts/group) and **(B)** representative infarct images, and **(C)** functional recovery in hearts from male and female hearts (**p<0.01 vs. male + formate, female; n = 5-7 hearts/group).

### Formate is protective in a model of in vitro I/R injury

We next sought to determine the mechanism of formate-induced protection, so we conducted *in vitro* experiments using H9c2 cells to evaluate formate-induced effects on protein SNO levels as a measure of NO production. Interestingly, formate-treated cells showed an increase in protein SNO levels compared to untreated cells after 48 hrs of treatment with 10 µmol/L formate (Fig. 3A). However, this formate-induced increase in protein SNO was blocked with L-NAME, supporting a role for NOS. To further probe mechanism, we next used an established *in vitro* model of I/R injury with NMVMs, as described previously.^29, 30^ To emulate our *ex vivo* findings, we treated NMVMs with 30 μmol/L formate for one hour prior to I/R injury. Importantly, formate conferred protection with this acute exposure period, significantly reducing LDH release compared to untreated cells (Fig. 3B). We next inhibited NOS activity with L-NAME and found that formate-induced protection was blocked (Fig. 3B). Collectively, these *in vitro* findings suggest that formate increases protein SNO through a NOS-dependent mechanism.

**Figure 3.**
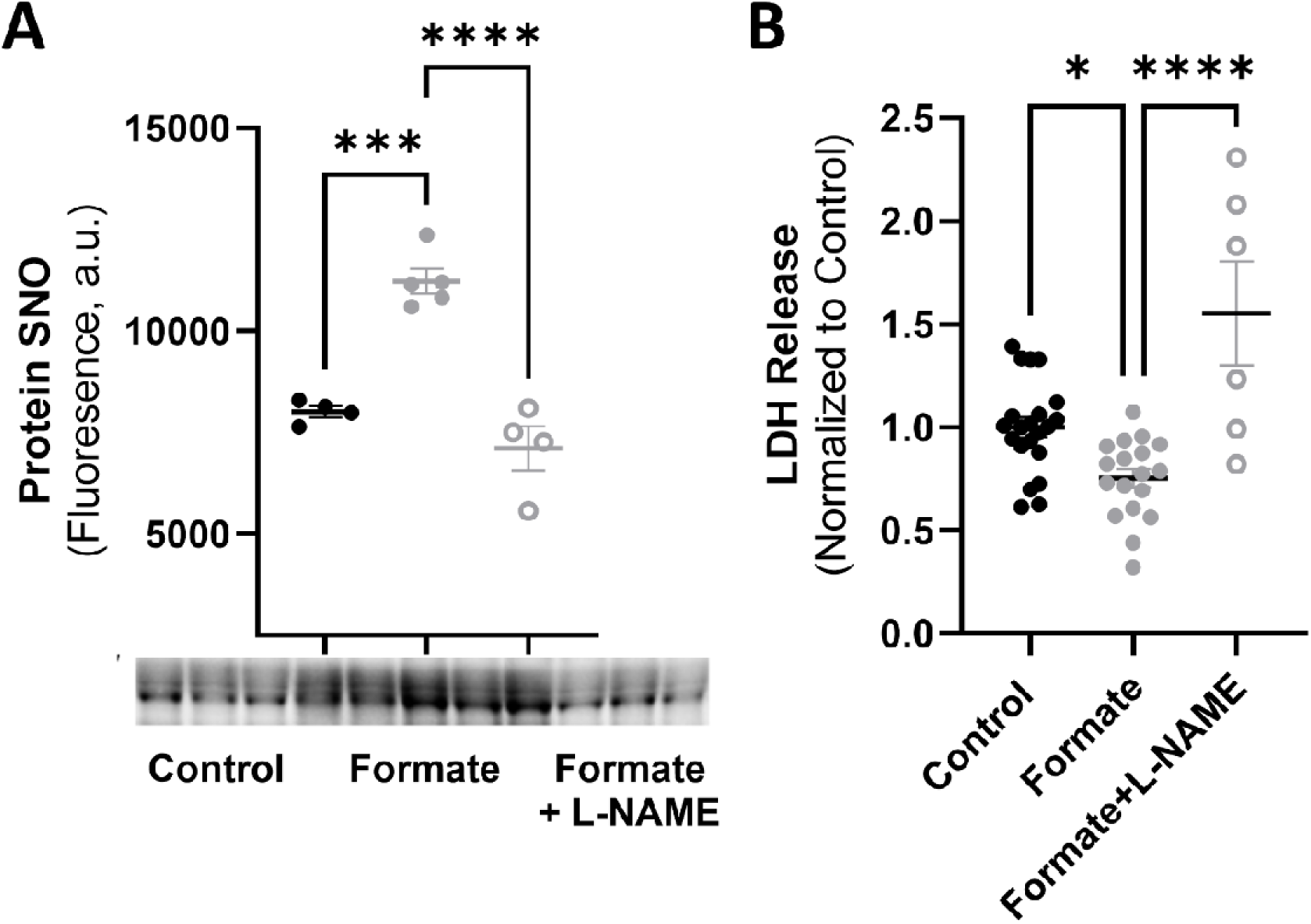
Exogenous formate reduces *in vitro* I/R injury in NMVMs. **(A)** Protein SNO levels from H9c2 cells treated with +/- 10 µmol/L formate and +/- 10 µmol/L L-NAME (non-specific NO synthase inhibitor) (***p<0.001 vs. formate, ****p<0.0001 vs. formate+L-NAME; n = 4-5 replicates/group). **(B)** NMVMs were treated with +/- 30 µmol/L formate and +/- 10 µmol/L L-NAME (non-specific NO synthase inhibitor) and subjected to coverslip-induced I/R injury. Lactate dehydrogenase (LDH) release in media as a measure of injury (*p<0.05 vs. formate, ****p<0.0001 vs. formate+L-NAME; n = 6-21 replicates/group).

### NOS inhibition blocks formate-induced cardioprotection

We further examined a potential role for NOS in formate-induced protection by moving back to our *ex vivo* model of I/R injury. Formate treatment significantly reduced infarct size in male hearts vs. untreated controls, while non-specific NOS inhibition with L-NAME attenuated this formate-mediated protection (Fig. 4). As such, NOS function appears to be a necessary component of formate-mediated protection from I/R injury.

**Figure 4.**
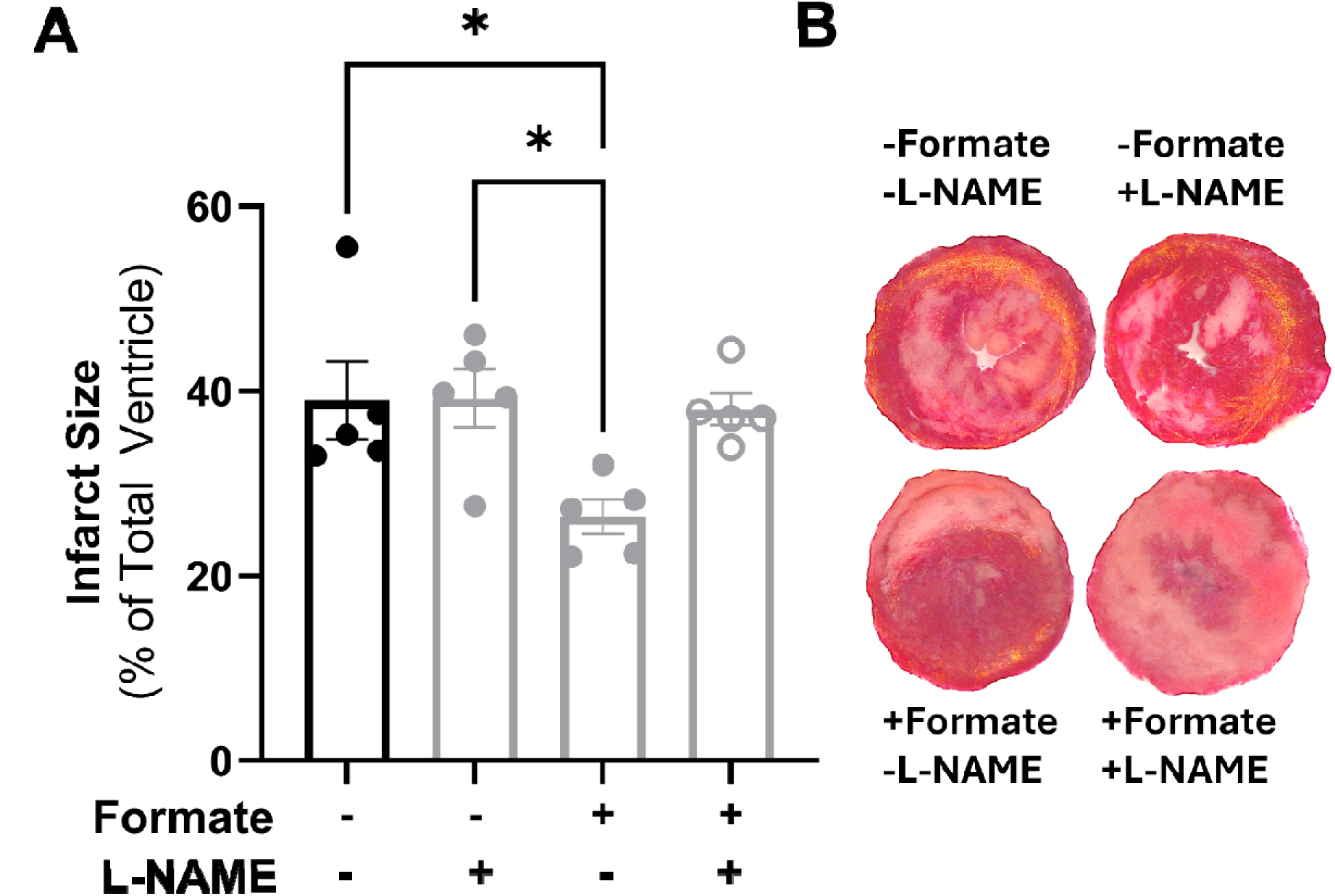
Formate-induced protection is blocked with NOS inhibition. Male hearts were Langendorff-perfused +/- 30 µmol/L formate and +/- 10 µmol/L L-NAME (non-specific NO synthase inhibitor), and subjected to I/R injury. **(A)** Infarct size (*p<0.05 vs. - formate/-L-NAME, -formate/+L-NAME, +formate/+L-NAME; n=5 hearts/group) and **(B)** representative infarct images from male hearts.

### Formate perfusion increases post-ischemic SNO levels

We also conducted additional *ex vivo* experiments to further validate our *in vitro* findings. Langendorff-perfused male hearts were perfused with or without 30 µmol/L formate and subjected to I/R injury. Post-reperfusion protein SNO levels were significantly increased in formate-perfused hearts at 2 mins (Fig. 5A) and 15 mins (Fig. 5B-C) of reperfusion compared to untreated hearts. These data suggest that formate contributes to cardioprotection by increasing protein SNO levels.

**Figure 5.**
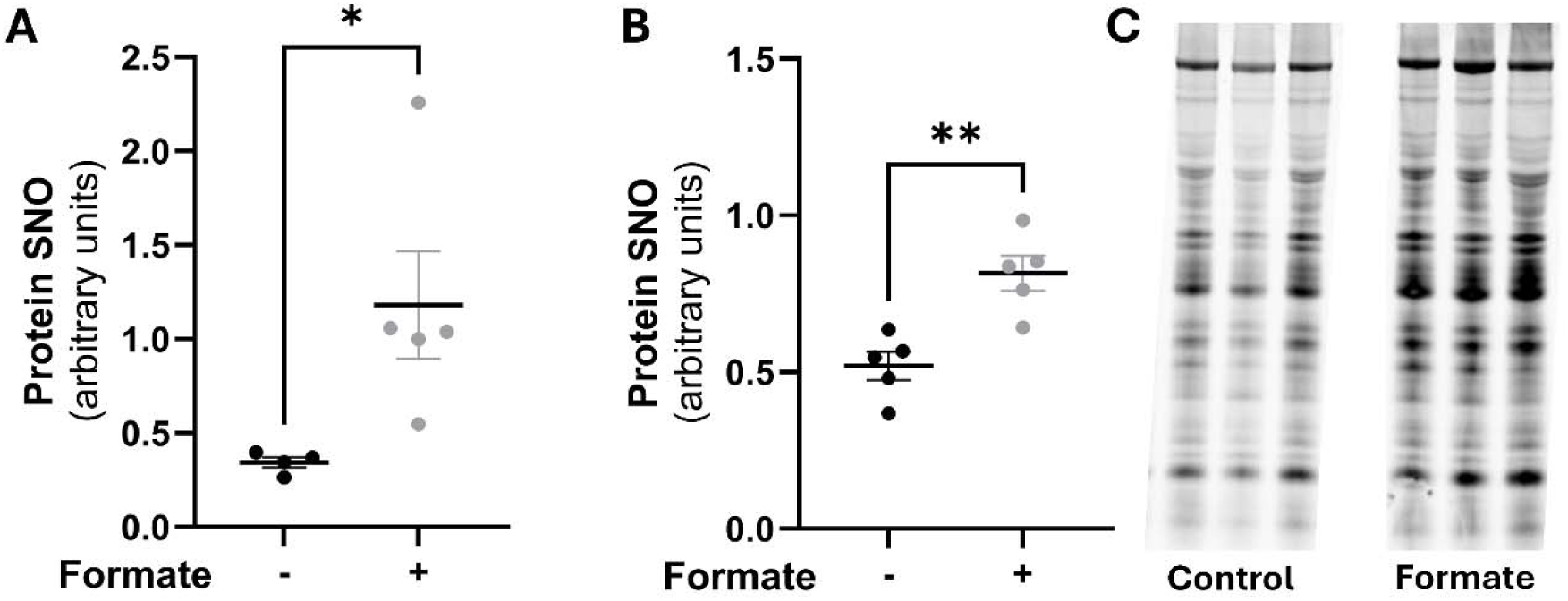
Formate enhanced post-ischemic SNO levels. Male hearts were Langendorff- perfused +/- 30 µmol/L formate, subjected to I/R injury and snap frozen after 2 and 15 mins of reperfusion. Protein SNO levels were measured via modified biotin switch at **(A)** 2 mins post-reperfusion (*p<0.05, n=4-5 hearts/group) and **(B)** 15 mins post-reperfusion (**p<0.01, n=5 hearts/group). (C) Representative image of SNO protein labeling.

### Formate increases ADH5 activity

To determine a potential mechanism for the formate-induced increase in protein SNO, we initially hypothesized that formate increases protein SNO by inhibiting ADH5 through a negative feedback loop,^36^ since formate can be produced via ADH5 activity. ADH5 activity was assessed in male hearts perfused with and without 30 µmol/L formate, in the presence or absence of 10 µmol/L L-NAME under non-ischemic conditions or following I/R injury, collected 2 mins after the onset of reperfusion. Simple linear regression revealed an overall trend towards significance across experimental groups. Subsequent ANOVA with multiple comparisons revealed that, under non-ischemic conditions, formate-treated hearts unexpectedly showed greater ADH5 activity than untreated controls (data not shown). I/R injury decreased ADH5 activity in both untreated and formate-treated hearts, though this decrease did not reach statistical significance in formate-treated hearts (Fig. 6). Importantly, formate-treated hearts subjected to I/R injury had significantly increased ADH5 activity as compared to nontreated hearts (Fig. 6). Co-administration of L-NAME attenuated the formate-mediated increase in ADH5 activity in ischemic hearts, suggesting that upstream NO signaling shapes formate-induced changes in ADH5 activity (Fig. 6). Together, these data demonstrate that formate increases ADH5 activity in both normal and postischemic conditions, suggesting that changes in ADH5 activity are unlikely to support a formate-mediated increase in protein SNO levels.

**Figure 6.**
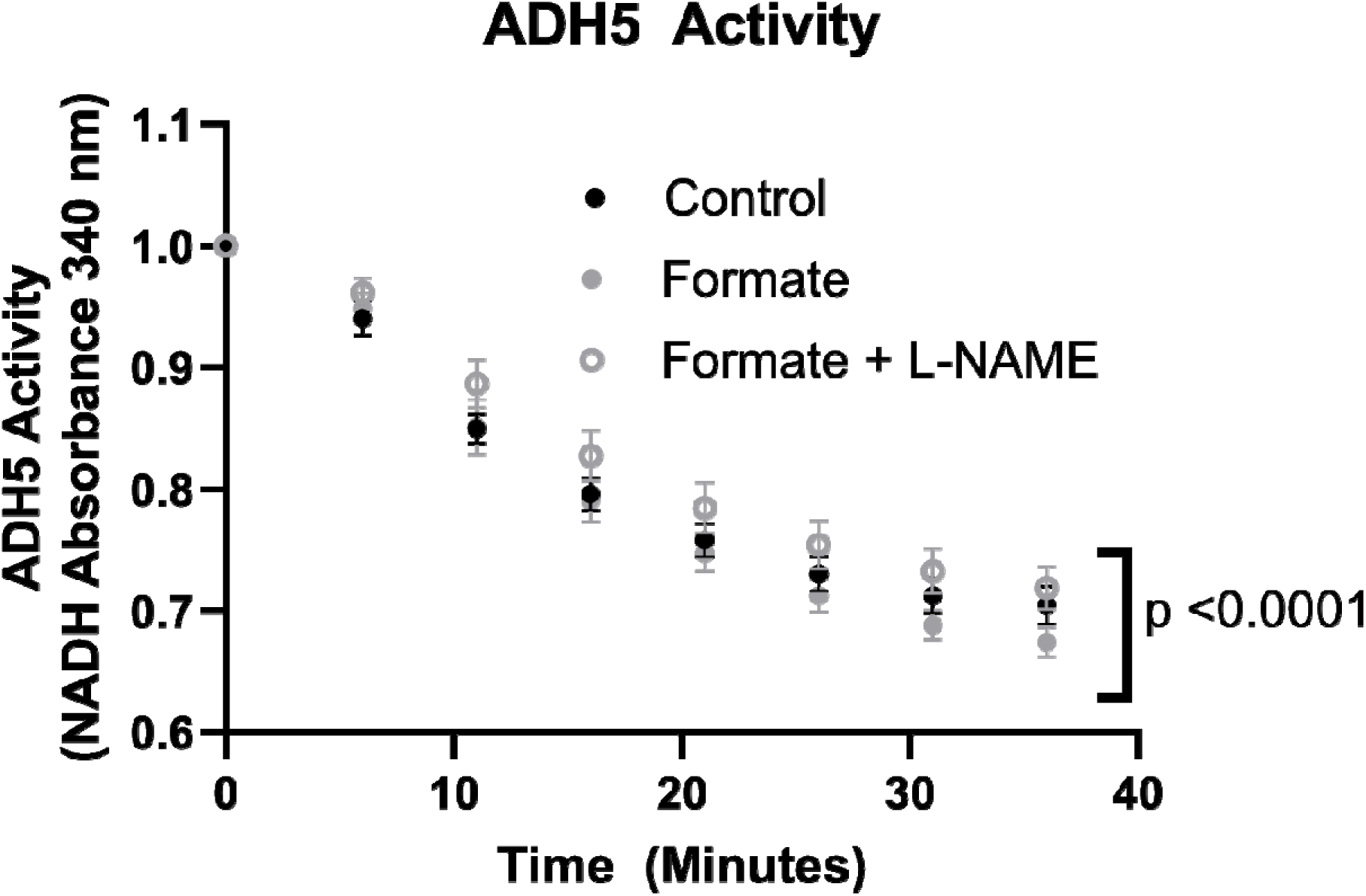
Formate increases ADH5 activity. Male hearts were Langendorff-perfused +/- 30 µmol/L formate and +/- 10 µmol/L L-NAME (non-specific NO synthase inhibitor), subjected to I/R injury and snap frozen after 2 mins of reperfusion. ADH5 activity via NADH-linked assay in male hearts (p<0.0001; n = 5 hearts/group).

### Formate preserves post-ischemic BH_4_ Levels

Formate is imported into the cell via the SLC26A6 transporter,^37^ which is expressed in the heart.^38^ Formate can then be used in one-carbon metabolism (OCM) to recycle the essential NOS co-factor BH_4_ through a salvage pathway. Therefore, we next examined an alternative hypothesis that formate alters BH_4_ levels. Male hearts were subjected to I/R injury with and without formate, and BH_4_, dihydrobiopterin (BH_2_), and total biopterin (B) levels were assessed in hearts at 2 mins of reperfusion (Fig. 7). Interestingly, untreated hearts showed a significant decrease in BH_4_ levels (Fig. 7A) and a corresponding increase in BH_2_ levels (Fig. 7B), likely due to the oxidation of BH_4_ to BH_2_. However, formate-treated hearts did not show a decrease in BH_4_, an increase in BH_2_, or a change in the BH_4_/BH_2_ ratio, indicating that BH_4_ levels are maintained. Furthermore, total biopterin levels did not differ among any of the groups. These data support a role for BH_4_ in formate-mediated cardioprotection.

**Figure 7.**
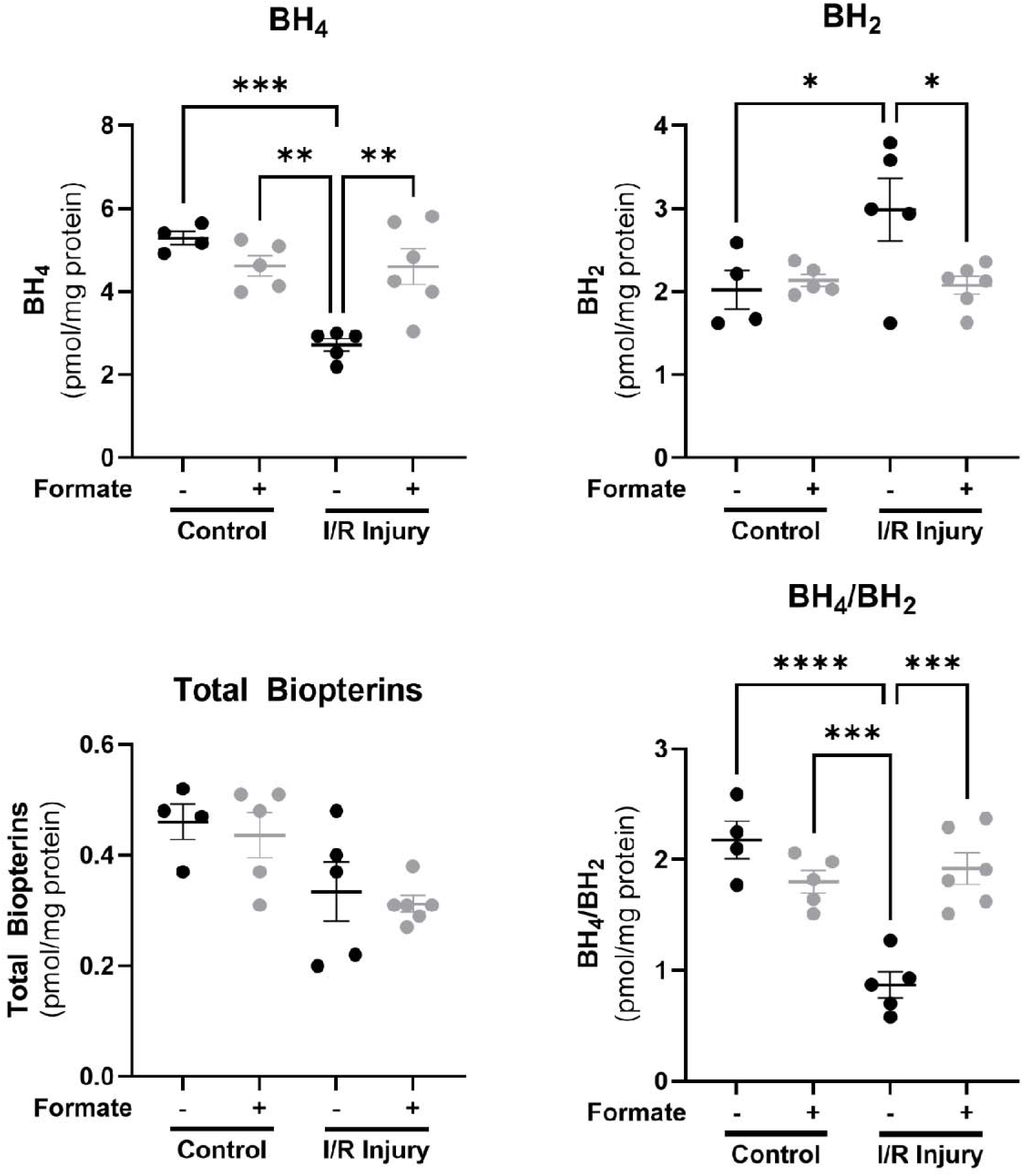
Formate preserves post-ischemic BH_4_ levels. Male hearts were Langendorff-perfused +/- 30 µmol/L formate, subjected to I/R injury and snap frozen after 2 mins of reperfusion. **(A)** BH_4_, **(B)** BH_2_, **(C)** total biopterin and **(D)** BH_4_/BH_2_ levels were measured via HPLC (**p<0.01, ***p<0.001, ****p<0.0001; n = 4-5 hearts/group).

### Inhibition of BH_4_ recycling blocks formate-induced protection

DHFR is the OCM enzyme that is responsible for recycling BH_4_ from BH_2_. Therefore, in a final set of experiments, to examine a potential role for OCM and DHFR, we inhibited DHFR activity using methotrexate. Importantly, we found a statistical trend towards blunted formate-mediated cardioprotection with methotrexate when examining infarct size in male hearts (Fig. 8) compared to hearts treated with formate alone. These findings suggest that DHFR and OCM may play a role in formate-mediated cardioprotection.

**Figure 8.**
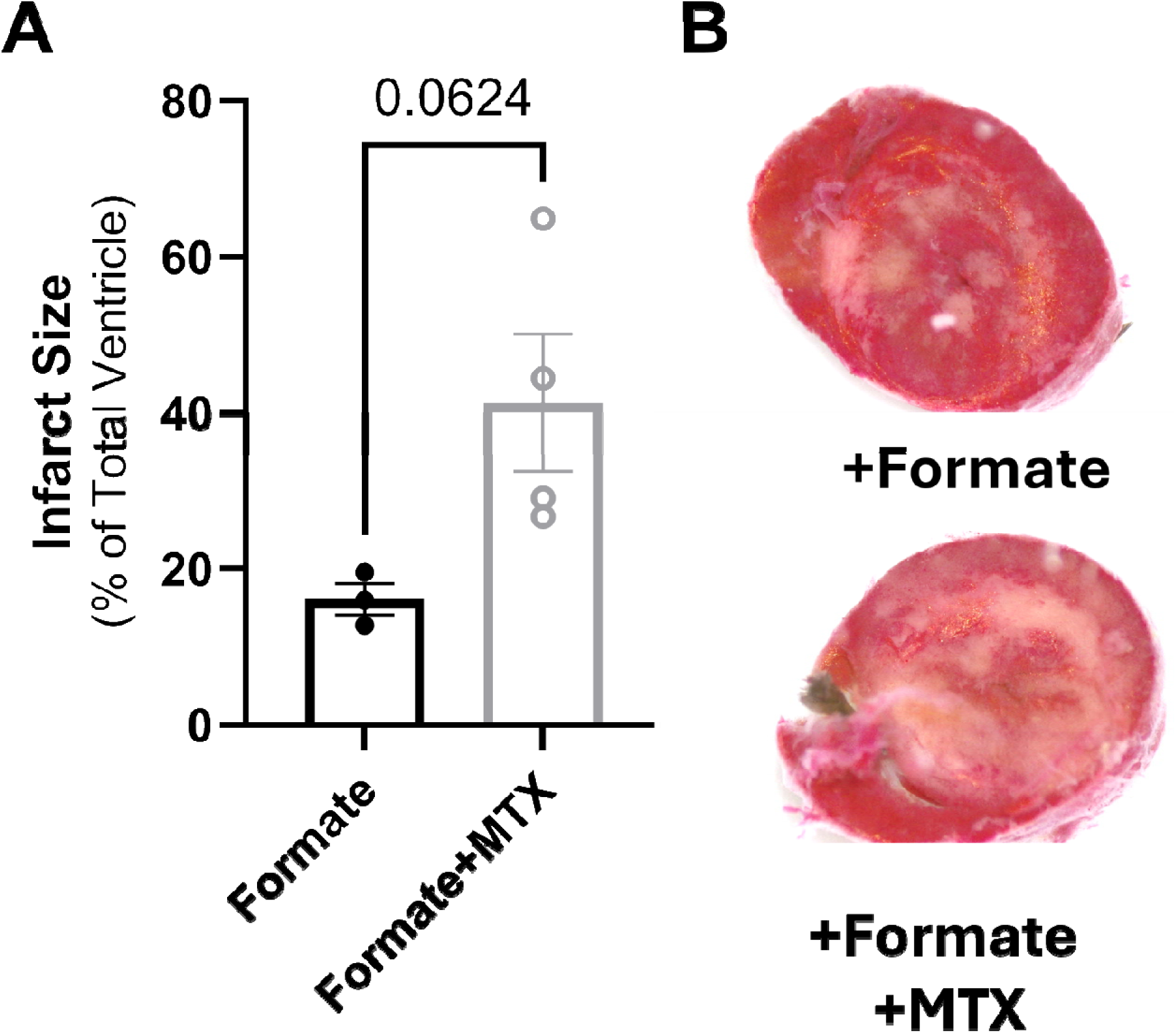
DHFR enzyme inhibition blunts formate-mediated protection in males. Male hearts were Langendorff-perfused +/- 30 µmol/L formate and +/- 10 µmol/L Methotrexate (MTX, DHFR inhibitor) and subjected to I/R injury. **(A)** Infarct size and **(B)** representative infarct images from male hearts (p = 0.0624; n=3-4 hearts/group).

## DISCUSSION

Herein, we demonstrate, for the first time, that formate substantially reduces ischemic injury in male hearts (Fig. 2) and explore the underlying protective mechanism(s). We find that formate confers robust protection from I/R injury in the male heart *ex vivo* (Fig. 2) and in NMVMs *in vitro* (Fig. 3). Mechanistically, formate-mediated protection increased protein SNO levels in both *in vitro* and *ex vivo* models (Figs. 3, 5), and protection was abrogated with non-specific NOS inhibition (Figs. 3, 4). Moreover, formate-mediated protection was associated with the preservation of BH_4_ levels (Fig. 7), and protection appeared blunted upon inhibition of the OCM enzyme DHFR (Fig. 8). Collectively, our findings support a cardioprotective mechanism whereby exogenous formate is used in OCM to facilitate the recycling of BH_4_ and increase protein SNO levels though a NOS-dependent mechanism.

Formate, or formic acid, was originally discovered as a noxious component of ant and bee venom, and while formate can be toxic at high levels like those associated with methanol toxicity, formate is produced endogenously in mammals via the gut microbiome and in cells through biochemical reactions that circulate in the range of 20-100 μmol/L in humans and other animals.^39–42^ Formate is imported into the cell via the SLC26A6 transporter,^37^ which shows high expression in the myocardium.^38^ Formate toxicity is further reduced via rapid assimilation into one-carbon metabolism (OCM), which is an interconnected network of biochemical reactions occurring in the cytosol and mitochondria that support numerous cellular functions, including epigenetic regulation, energetics, and the maintenance of redox homeostasis.^43^ Recent studies suggest that ADH5 and ALDH2 contribute to circulating formate levels,^44^ and our prior work has shown that ADH5 activity is reduced in male hearts compared to females.^12, 20, 21^ Our current findings show that ALDH2 activity is similarly reduced in male hearts (Fig. 1). These findings are consistent with a prior study demonstrating reduced ALDH2 phosphorylation and activity in male rat hearts compared to females.^35^ As such, we surmised that endogenous formate production may be reduced in male hearts, and exogenous formate administration may yield cardioprotective benefits. Indeed, we found that the administration of exogenous formate yielded substantial protection from ischemic injury in males (Fig. 2). Exogenous formate also reduced injury in our established in vitro model of coverslip-induced ischemia with NMVMs (Fig. 3).

Furthermore, as we have shown in previous studies,^12, 20–22^ female hearts exhibited a sex-specific reduction in ischemic injury at baseline compared to males (Fig. 2), but formate did not confer additional protective benefit (Fig. 2). As mentioned, we have found that compared to males, female hearts exhibit higher ADH5 and ALDH2 activity, suggesting that endogenous circulating formate levels may be increased. In support of this idea, plasma formate is higher in pre-menopausal women compared to age-matched men.^40^ Female mice also have higher plasma formate than males,^45^ and whole body formate consumption appears to differ by sex.^45^ As such, supplementation with additional exogenous formate may be redundant for female hearts, thereby conferring no additional protective benefit. Collectively, our findings reveal formate as a novel mediator of cardioprotective signaling in male hearts, and therefore we further examined underlying protective mechanism(s).

### Nitric oxide and protein SNO

Our prior work supports protein SNO as a critical stress response to ischemia that yields cardioprotective benefit by maintaining redox homeostasis. In particular, we and others have shown that cardioprotective interventions which increase protein SNO, strongly correlate with reduced oxidative stress and ischemic injury, particularly in male hearts.^6–14^ Since myocardial SNO protein homeostasis is mediated in part via NOS-dependent activity, we examined the impact of non-specific NOS inhibition on formate mediated protection *in vitro* and *ex vivo* in male hearts, and found that L-NAME alleviated the protection afforded by formate in both models (Figs. 3, 4). These findings support NOS as a target of formate, and are consistent with prior studies that have consistently shown a protective role for NOS in myocardial ischemic injury. Moreover, we found that formate also increased protein SNO levels both *in vitro* (Fig. 3) and *ex vivo* (Fig. 5), providing further support for a protective role for the protein SNO in myocardial ischemic injury. Protein SNO is a reversible modification that can alter protein function, localization, and activity.^46^ We have also shown that protein SNO can shield cysteine residues from oxidative damage,^6, 8, 47^ which is critical in the context of ischemic injury. Recent work from our group and others has shown that SNO protein levels increase in male hearts with many different forms of cardioprotective stimuli, ranging from ischemic pre- and post-conditioning to the use of pharmacologic agents.^6–11, 13, 14, 48, 49^ These protective effects can be blocked with the SNO-specific reducing agent ascorbate,^10^ but not with soluble guanylate cyclase or PKG inhibition.^9, 49–51^ As such, our current findings are consistent with our prior work showing that interventions that increase protein SNO levels are protective in male hearts, but NOS inhibition alleviates beneficial effects.^11,43^

### One-carbon metabolism and BH_4_

Formate can be imported into OCM via methylenetetrahydrofolate dehydrogenase 1/2 (MTHFD1/2) and used as a one-carbon donor to maintain redox homeostasis by contributing to the synthesis of antioxidants that include NADPH and glutathione.^45, 52^ Formate can also be used by OCM to drive the salvage and recycling of the critical NOS cofactor BH_4_ from its oxidized form (BH_2_) by the OCM enzyme DHFR.^40^ Notably, in the current study, untreated hearts subjected to I/R injury showed a significant decrease in BH_4_ levels and a significant increase in BH_2_. However, while BH_4_ and BH_2_ levels remained unchanged with formate treatment under non-ischemic conditions, formate significantly preserved BH_4_ levels, reduced BH_2_ levels and improved the BH_4_/BH_2_ ratio following I/R injury (Fig. 7). Additionally, total biopterin levels did not differ among groups with or without formate treatment or I/R injury (Fig. 7). Since BH_4_ can be generated via *de novo* synthesis through a enzymatic pathway that includes guanosine triphosphate (GTP) and GTP cyclohydrolase I, and through a separate OCM-dependent salvage pathway,^53^ our findings support a formate-mediated increase BH_4_ recycling, as opposed to *de novo* synthesis. DHFR is the OCM enzyme that plays a key role in recycling BH_4_ from BH_2_, and we have previously shown that this is a key process in the maintenance of endothelial NOS (eNOS) coupling and function.^34^ Importantly, inhibition of DHFR with methotrexate appeared to blunt formate-mediated protection in the current study (Fig. 8), suggesting a potential role for DHFR in formate-mediated protection, but these data did not reach statistical significance. Although a link between formate and BH_4_ recycling has not previously been established, the one-carbon donor folate has been shown to increase BH_4_ levels and eNOS function in endothelial cells *in vitro* through a DHFR-dependent mechanism,^54^ and to lessen ischemic heart injury *in vivo* by reducing eNOS uncoupling.^55, 56^ The OCM intermediate 5-methyl-tetrahydrofolate (5-MTHF) is thought to be involved,^57, 58^ as 5-MTHF improves eNOS function by increasing BH_4_ bioavailability.^57, 58^ 5-MTHF has also been shown to restore NO production in patients with heart disease.^59^ Importantly, formate can be converted to 5-MTHF by a series of OCM enzymes. Collectively, our findings suggest that formate-mediated cardioprotection occurs through a DHFR-dependent mechanism that preserves BH_4_ levels, but additional studies are necessary to confirm this mechanism.

### Study Limitations

There are a number of limitations in this study that require acknowledgement. Firstly, formate can act directly as an antioxidant,^60^ and while our findings suggest that formate mediates BH_4_ recycling, we cannot rule out potential antioxidant contributions. Additionally, only one dose of formate was evaluated, while plasma formate circulates in the range of 20-100 μmol/L in mammals.^39–42^ The timing of formate administration was also not fully evaluated, as we administered formate for 20 mins before ischemia and during the entirety of reperfusion. As such, future studies will need to examine additional dose- and time-dependent effects of exogenous formate. Furthermore, we only examined one dose of methotrexate in male hearts, so future studies will need to examine the dose-dependent effects of methotrexate as well.

### Conclusions

In conclusion, the results of our study support critical roles for BH_4_, NOS and protein SNO, as well as DHFR, in formate-mediated cardioprotection. Specifically, we found that formate preserved post-ischemic BH_4_ levels, and increased protein SNO levels through a NOS-dependent mechanism. Furthermore, we found that DHFR inhibition appeared to blunt formate-mediated protection, suggesting that there may be a potential link between OCM and NO signaling in formate-mediated protection. Collectively, our findings identify formate utilization by OCM as a novel pathway to increase protein SNO and drive protection from ischemic injury in the male heart. These findings represent an important step towards the treatment of ischemic heart disease in males.

## GRANTS

This work was supported by the National Institutes of Health [T32 ES007141 (HG), F31 HL165820 (HG), R21 HL157800 (MK) and R01 HL136496 (MK)] and the American Heart Association [TPA 970850 (SD) and 23DIVSUP1057308 (SD)].

## AUTHOR CONTRIBUTIONS

H.G. and M.J.K conceived and designed research; H.G-G., R.K., O.V.E., P.P., D.A., D.Q., and M.C. performed experiments; H.G-G., R.K., O.V.E., P.P., D.A., D.Q., and M.C. analyzed data; H.G-G. and M.J.K. interpreted results of experiments; H.G-G. and M.J.K. prepared figures; H.G-G. and M.J.K. drafted manuscript; H.G-G., R.K., O.V.E., P.P., D.A., D.Q., S.D., B.O., M.C. and M.J.K. edited and revised the manuscript; H.G-G., R.K., O.V.E., P.P., D.A., D.Q., S.D., B.O., M.C. and M.J.K. approved the final version of this manuscript.

## DATA AVAILABILITY STATEMENT

**Included in article.** The data that support the findings of this study are available in the Materials and Methods, Results, and/or Figures of this article.

